# Whole genome sequencing, *de novo* assembly and phenotypic profiling for the new budding yeast species *Saccharomyces jurei*

**DOI:** 10.1101/339929

**Authors:** Samina Naseeb, Haya Alsammar, Tim Burgis, Ian Donaldson, Norman Knyazev, Christopher Knight, Daniela Delneri

**Author notes:** These authors contributed equally in this work (Samina Naseeb, Haya Alsammar). Corresponding authors: Samina Naseeb Daniela Delneri.

## Abstract

*Saccharomyces sensu stricto* complex consist of yeast species, which are not only important in the fermentation industry but are also model systems for genomic and ecological analysis. Here, we present the complete genome assemblies of *Saccharomyces jurei,* a newly discovered *Saccharomyces sensu stricto* species from high altitude oaks. Phylogenetic and phenotypic analysis revealed that *S. jurei* is a sister-species to *S. mikatae,* than *S. cerevisiae,* and *S. paradoxus.* The karyotype of *S. jurei* presents two reciprocal chromosomal translocations between chromosome VI/VII and I/XIII when compared to *S. cerevisiae* genome. Interestingly, while the rearrangement I/XIII is unique to *S. jurei,* the other is in common with *S. mikatae* strain IFO1815, suggesting shared evolutionary history of this species after the split between *S. cerevisiae* and *S. mikatae.* The number of Ty elements differed in the new species, with a higher number of Ty elements present in *S. jurei* than in *S. cerevisiae.* Phenotypically, the *S. jurei* strain NCYC 3962 has relatively higher fitness than the other strain NCYC 3947^T^ under most of the environmental stress conditions tested and showed remarkably increased fitness in higher concentration of acetic acid compared to the other *sensu stricto* species. Both strains were found to be better adapted to lower temperatures compared to *S. cerevisiae.*

## Introduction

*Saccharomyces sensu stricto* yeasts, currently comprise eight species: S. *cerevisiae, S. paradoxus, S. uvarum, S. mikatae, S. kudriavzevii, S. arboricola, S. eubayanus, S. jurei* (Libkind et al. 2011; Martini and Martini 1987; Naseeb et al. 2017b; Naumov et al. 1995.a; Naumov et al. 1995. b; Naumov et al. 2000; Wang and Bai 2008) and two natural hybrids: *S. pastorianus* (Masneuf et al. 1998; Querol and Bond 2009) and *S. bayanus* (Nguyen et al. 2011). *Saccharomyces jurei* is the latest addition to the *sensu stricto* clade and was isolated from oak tree bark and surrounding soil at an altitude of 1000m above sea level in Saint Auban, France (Naseeb et al. 2017b). It is known that species within *sensu stricto* group are reproductively isolated and possess post-zygotic barriers (Naumov 1987). Moreover, yeasts within this group exhibit almost identical karyotypes with 16 chromosomes (Cardinali and Martini 1994; Carle and Olson 1985; Naumov et al. 1996).

In the modern era of yeast genetics, the advances in sequencing technology has lead to the whole genome sequencing of many *Saccharomyces sensu stricto* species (*S. cerevisiae*, *S. bayanus var. uvarum, S. kudriavzevii, S. mikatae, S. paradoxus, S. eubayanus* and *S. arboricola)* (Casaregola et al. 2000; Cliften et al. 2003; Kellis et al. 2003; Libkind et al. 2011; Liti et al. 2013; Scannell et al. 2011). To date, more than 1000 *S. cerevisiae* strains belonging to different geographical and environmental origins have been sequenced and assembled (Engel and Cherry 2013; Peter et al. 2018a). The availability of sequencing data from multiple strains of hemiascomycets yeast species has enhanced our understanding of biological mechanisms and comparative genomics. Researchers are now combining comparative genomics with population ecology to better understand the genetic variations, taxonomy, evolution and speciation of yeast strains in nature. Genome variation provides the raw material for evolution, and may arise by various mechanisms including gene duplication, horizontal gene transfer, hybridization and micro and macro rearrangements (Fischer et al. 2001; Hall et al. 2005; Lynch 2002; Naseeb and Delneri 2012; Naseeb et al. 2016; Naseeb et al. 2017a; Seoighe et al. 2000). Synteny conservation studies have shown highly variable rates of genetic rearrangements between individual lineages both in vertebrates and in yeasts (Bourque et al. 2005; Fischer et al. 2006). This genome variation is a means of evolutionary adaptation to environmental changes. An understanding of the genetic machinery linked to phenotypic variation provides knowledge of the distribution of *Saccharomyces* species in different environments, and their ability to withstand specific conditions (Brice et al. 2018; Goddard and Greig 2015; Jouhten et al. 2016; Peter et al. 2018b).

Recently, we isolated two strains (NCYC 3947^T^ and NCYC 3962) of *Saccharomyces jurei* from *Quercus robur* bark and surrounding soil (Naseeb et al. 2017b). The initial sequencing of ITS1, D1/D2 and seven other nuclear genes showed that both strains of *S. jurei* were closely related to *S. mikatae* and *S. paradoxus* and grouped in *Saccharomyces sensu stricto* complex. We also showed that *S. jurei* can readily hybridize with other *sensu stricto* species but the resulting hybrids were sterile (Naseeb et al. 2017b). Here, we represent high quality *de novo* sequence and assembly of both strains (NCYC 3947^T^ and NCYC 3962) of *S. jurei.* The phylogenetic analysis placed *S. jurei* in the *sensu stricto* clade, in a small monophyletic group with *S. mikatae.* By combining Illumina HiSeq and PacBio data, we were able to assemble full chromosomes and carry out synteny analysis. Moreover, we show that *S. jurei* NCYC 3962 had higher fitness compared to NCYC 3947^T^ under different environmental conditions. Fitness of *S. jurei* strains at different temperatures showed that it was able to grow at wider range of temperatures (12°C-37°C).

## Material and Methods

### Yeast strains

Strains used in this study are presented in Table 1. All strains were grown and maintained on YPDA (1% w/v yeast extract, 2% w/v Bacto-peptone, 2% v/v glucose and 2% w/v agar). Species names and strains number are stated in Table 1.

**Table 1:** Strains used in this study.

| **Species** | **Strain number** | **References** |
| --- | --- | --- |
| *S. jurei* | NCYC 3947<sup>T</sup><br>NCYC 3962 | (Naseeb et al. 2017b) |
| *S. cerevisiae* | NCYC 505<sup>T</sup> | (Vaughan Martini and Kurtzman 1985) |
| *S. paradoxus* | CBS 432<sup>T</sup> | (Kaneko and Banno 1991) |
| *S. mikatae* | NCYC 2888<sup>T</sup> (IFO 1815<sup>T</sup>) | (Yamada et al. 1993) |
| *S. kudriavzevii* | NCYC 2889<sup>T</sup> (IFO 1802<sup>T</sup>) | (Yamada et al. 1993) |
| *S. arboricola* | CBS 10644<sup>T</sup> | (Wang and Bai 2008) |
| *S. eubayanus* | PYCC 6148<sup>T</sup> (CBS 12357<sup>T</sup>) | (Libkind et al. 2011) |
| *S. uvarum* | NCYC 2669 (CBS 7001<sup>T</sup>) | (Pulvirenti et al. 2000) |
| *S. pastorianus* | NCYC 329<sup>T</sup> (CBS 1538<sup>T</sup>) | (Martini and Martini 1987) |

### DNA Extraction

For Illumina Hiseq, the total DNA was extracted from an overnight grown culture of yeast strains by using the standard phenol/chloroform method described previously (Fujita and Hashimoto 2000) with some modifications. Briefly, 5 ml of overnight grown yeast cells were centrifuged and resuspended in 500 μl EB buffer (4M sorbitol, 500mM EDTA and1M DTT) containing 1 mg/ml lyticase. The cells were incubated at 37°C for 1 hour. Following incubation, the cells were mixed with stop solution (3M NaCl, 100mM Tris pH 7.5 and 20mM EDTA) and 60 μl of 10% SDS. The cell suspension was vortexed and mixed with 500 μl phenol-chloroform. The samples were centrifuged at 13000 rpm for 2 minutes to separate the aqueous phase from the organic phase. The upper aqueous phase was transferred to a clean 1.5 ml tube and phenol-chloroform step was repeated twice until a white interface was no longer present. The aqueous phase was washed with 1 ml absolute ethanol by centrifugation at 13000 rpm for 10 minutes. The pellet was air dried and resuspended in 30 μl of sterile milliQ water.

Genomic DNA for PacBio sequencing was extracted using Qiagen Genomic-tip 20/G kit (cat. No. 10223) following manufacturer’s recommended instructions. The yield of all DNA samples was assessed by the nanodrop spectrophotometer (ND-1000) and by Qubit 2.0 fluorometer (catalogue no. Q32866). Purity and integrity of DNA was checked by electrophoresis on 0.8% (w/v) agarose gel and by calculating the A260/A280 ratios.

### Library preparation for Illumina and PacBio sequencing

Paired end whole-genome sequencing was performed using the Illumina HiSeq platform. FastQC (Babraham Bioinformatics) was used to apply quality control to sequence reads, alignment of the reads was performed using BOWTIE2 (Langmead and Salzberg 2012) and post-processed using SAMTOOLS (Li et al. 2009).

For Pacbio sequencing, genomic DNA (10 μg) of NCYC 3947^T^ and NCYC 3962 strains was first DNA damage repaired, sheared with Covaris G-tube, end repaired and exonuclease treated. Smrtbell library (10-20kb size) was prepared by ligation of hairpin adaptors at both ends according to PacBio recommended procedure (Pacific Bioscience, No: 100-259-100). The resulting library was then size selected using Blue Pippin with 7-10kb cutoff. Sequencing run was performed on PacBio RS II using P6/C4 chemistry for 4 hours. The genome was assembled using SMRT analysis and HGAP3 pipeline was made using default settings.

### Genome assembly, annotation, orthology and chromosomal structural plots

The PacBio sequences were assembled using hierarchical genome-assembly process (HGAP) (Chin et al. 2013). Protein coding gene models were predicted using Augustus (Stanke and Morgenstern 2005) and the Yeast Genome Annotation Pipeline (Byrne and Wolfe 2005). In addition, protein sequences from other *Saccharomyces* species were aligned to the genome assembly using tblastn (Gertz et al. 2006). These predictions and alignments were used to produce a final set of annotated genes with the Apollo annotation tool (Lewis et al. 2002). The protein sequences were functionally annotated using InterproScan (Jones et al. 2014). Orthologous relationships with *S. cerevisiae* S288C sequences were calculated using InParanoid (Berglund et al. 2008). Non-coding RNAs were annotated by searching the RFAM database (Nawrocki et al. 2015) using Infernal (Nawrocki and Eddy 2013). Further tRNA predictions were produced using tRNAscan (Lowe and Eddy 1997). Repeat sequences were identified in Repbase (Bao et al. 2015) using Repeat Masker (Smit et al. 2013–2015).

The dotplots were constructed by aligning *S. jurei* genome to the *S. cerevisiae* S288C genome using NUCmer and plotted using MUMmerplot (Kurtz et al. 2004). These features are available to browse via a UCSC genome browser (Kent et al. 2002) track hub (Raney et al. 2014). Single nucleotide polymorphisms (SNPs) were identified using Atlas-SNP2(Challis et al. 2012).

### Phenotypic assays

#### Temperature tolerance

Fitness of *S. jurei* strains and *Saccharomyces sensu stricto* type strains was examined using FLUOstar optima microplate reader at 12^°^C, 16^°^C, 20^°^C, 25^°^C, 30^°^C and 37^°^C. Cells were grown from a starting optical density (OD) of 0.15 to stationary phase in YPD (1% w/v yeast extract, 2% w/v Bacto-peptone and 2% w/v glucose) medium. The growth OD_595_ was measured every 5 minutes with 1 minute shaking for 72 hours. Growth parameters, lag phase (*λ*), maximum growth rate (μ_max_), and maximum biomass (*A*_max_) were estimated using R shiny app on growth curve analysis (https://kobchai-shinyapps01.shinyapps.io/growth_curve_analysis/).

#### Environmental stress

Strains were screened for tolerance to environmental stressors using a high-throughput spot assay method. Cells were grown in a 96-well plate containing 100 μl YPD in four replicates at 30°C for 48 hours. The yeast strains grown in 96-well plate were sub-cultured to a 384 well plate to achieve 16 replicates of each strain and grown at 30°C for 48 hours. Singer ROTOR ©HAD robot (Singer Instruments, UK) was used to spot the strains on (i) YPDA + 0.4% & 0.6% acetic acid, (ii) YPDA+ 4mM & 6mM H_2_O_2_, (iii) YPDA+ 2.5mM & 5mM CuSO4, (iv) YPDA+ 2% & 5% NaCl, (v) YPDA+ 5% & 10% Ethanol (vi) YPA+ 15% maltose and (vii) YPA+ 30% & 35% glucose. The spot assay plates were incubated at 30°C and high-resolution images of phenotypic plates were taken using phenobooth after 3 days of incubation (Singer Instruments, UK). The colony sizes were calculated in pixels using phenosuite software (Singer Instruments, UK) and the heat maps of the phenotypic behaviors were constructed using R shiny app (https://kobchai-shinyapps01.shinyapps.io/heatmap_construction/).

#### Data and reagent availability

Strains are available upon request. Supplemental files are available at FigShare (https://figshare.com/s/60bbbc1e98886077182a). Figure S1 shows alignment of the amino acid sequences of *MEL1* gene belonging to *S. jurei* NCYC 3947^T^ (Sj) and *S. mikatae* IFO 1816 (Sm). Table S1, Table S2, Table S3 and Table S4 list the genes, which are present in simple one to one orthologous relationship, in many to many relationship, in many to one relationship and in one to many relationship, respectively. Table S5 lists the genes that are present in *S. cerevisiae* but absent in *S. jurei*. Table S6 lists the genes which are present in *S. jurei* but absent in *S. cerevisiae.* Table S7 lists the genes which are used to construct the phylogenetic tree. Table S8 lists the genes which are potentially introgressed in *S. jurei* genome from *S. paradoxus.* Table S9, Table S10 and Table S11 show lag phase time (λ), maximum growth rate (*μ*max) and maximum biomass (*A*_max_) of *Saccharomyces species* used in this study, respectively.

## Results and Discussion

### High quality *de novo* sequencing and assembly of *S. jurei* genome

Genome sequencing of the diploid *S. jurei* NCYC 3947^T^ and NCYC 3962 yeast strains was performed using Illumina Hiseq and Pacbio platforms. We obtained approximately 9.02 × 10^5^ and 4.5 × 10^5^ reads for NCYC 3947^T^ and NCYC 3962 respectively. We obtained 2 × 101 bp reads derived from ~200 bp paired-end reads which were assembled in 12 Mb genome resulting in a total coverage of 250x based on high quality reads. The sequencing results and assembled contigs are summarized in Tables 2–4. By combining the Illumina mate pair and Pacbio sequencing we were able to assemble full chromosomes of *S. jurei* NCYC 3947^T^ and NCYC 3962 (Tables 5 and 6). The total genome size (~12 Mb) obtained for both strains of *S. jurei* was comparable to the previously published genomes of *Saccharomyces sensu stricto* species (Baker et al. 2015; Goffeau et al. 1996; Liti et al. 2013; Scannell et al. 2011).

**Table 2:**
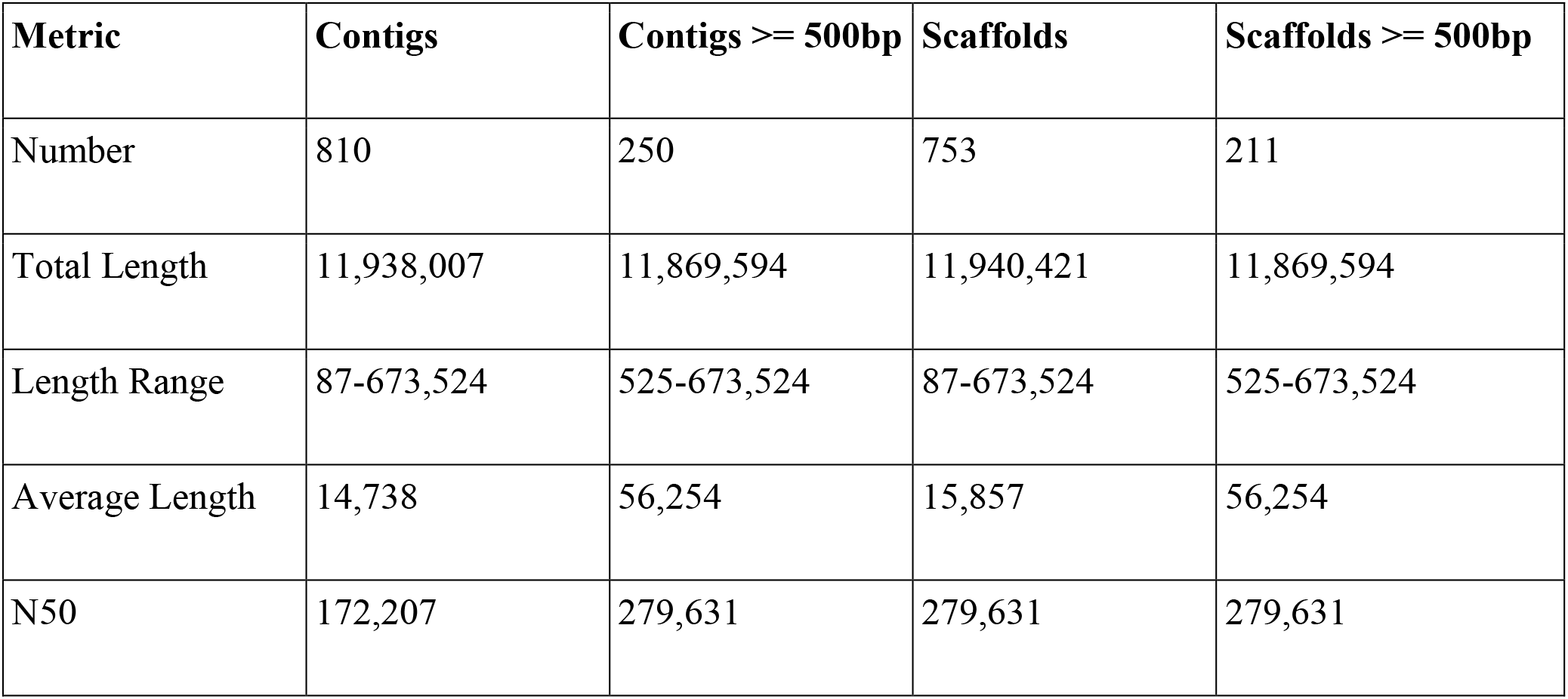
Summary of *S. jurei* NCYC 3947^T^ genome sequencing and assembly using Hi-seq platform.

**Table 3:** Summary of *S. jurei* NCYC 3962 genome sequencing and assembly using Hi-seq platform.

| **Metric** | **Contigs** | **Contigs >= 500bp** | **Scaffolds** | **Scaffold >= 500bp** |
| --- | --- | --- | --- | --- |
| Number | 3719 | 987 | 3618 | 933 |
| Total length | 11,760,925 | 11,419,281 | 11,768,034 | 11,441,494 |
| Length range | 59-80,684 | 507-80,684 | 59-80,684 | 507-80,684 |
| Average length | 3,162 | 11,569 | 3,252 | 12,263 |
| N50 | 20,806 | 21,318 | 21,928 | 22,552 |

**Table 4:** Summary of *S. jurei* NCYC 3947^T^ and NCYC 3962 genome assembly using PacBio platform.

| **Metric** | ***S. jurei* NCYC 3947** | ***S. jurei* NCYC 3962** |
| --- | --- | --- |
| Contigs | 35 | 57 |
| Max contig length | 1,474,466 | 1,470,125 |
| Contig N50 | 738,741 | 652,030 |
| Total assembly size | 12,306,756 | 12,932,708 |

**Table 5:** Total number of chromosomes assembled in *S. jurei* NCYC 3947^T^.

| **Sequence name** | **Length (bp) including gaps** |
| --- | --- |
| chrI.1_chrXIII.2 | 809,572 |
| chrII | 809,280 |
| chrIII | 308,350 |
| chrIV | 1,474,466 |
| chrV | 584,553 |
| chrVI.1_chrVII.2 | 730,011 |
| chrVI.2_chrVII.1 | 638,210 |
| chrVIII | 534,462 |
| chrIX | 434,517 |
| chrX | 738,741 |
| chrXI | 671,067 |
| chrXII.1 | 458,950 |
| chrXII.2 | 568,540 |
| chrI.2_chrXIII.1 | 334,136 |
| chrXIV | 749,072 |
| chrXV | 1,068,672 |
| chrXVI | 920,427 |
| chrMT | 105,732 |

**Table 6:** Total number of chromosomes assembled in *S. jurei* NCYC 3962.

| **Sequence name** | **Length (bp) including gaps** |
| --- | --- |
| chrI.1_chrXIII.2 | 756,315 |
| chrII | 814,183 |
| chrIII | 329,028 |
| chrIV | 1,470,125 |
| chrV | 570,437 |
| chrVI.1_chrVII.2 | 723,619 |
| chrVII.2_chrVI.1 | 652,030 |
| chrVIII | 536,516 |
| chrIX | 439,662 |
| chrX.1 | 487,336 |
| chrX.2 | 258,684 |
| chrXI | 676,065 |
| chrXII.1 | 475,978 |
| chrXII.2 | 571,082 |
| chrI.2_chrXIII.1 | 334,998 |
| chrXIV | 790,124 |
| chrXV.1 | 474,048 |
| chrXV.2 | 240,703 |
| chrXV.3 | 236,823 |
| chrXV.4 | 114,889 |
| chrXVI | 806,586 |
| chrMT | 110,829 |

### *S. jurei* genome prediction and annotation

The high-quality de novo assembly of *S. jurei* NCYC 3947^T^ genome resulted in 5,794 predicted protein-coding genes for *S. jurei,* which is similar to the published genomes of other *sensu stricto* species (Baker et al. 2015; Liti et al. 2009; Liti et al. 2013; Scannell et al. 2011; Walther et al. 2014). Of the predicted protein-coding genes, 5,124 were in a simple 1:1 putatively orthologous relationship between *S. cerevisiae* and *S. jurei* (Table S1). From the remaining protein-coding genes, 35 genes showed many to many relationship (multiple *S. cerevisiae* genes in paralogous cluster with multiple *S. jurei* genes (Table S2), 31 genes were in many to one relationship (many genes in *S. cerevisiae* are in an paralogous cluster with a single *S. jurei* gene; most of these were found to be retrotransposons; Table S3) and 50 genes were in one to many relationships (one *S. cerevisiae* gene in an paralogous cluster with many *S. jurei* genes; Table S4). Interestingly, we found an increase in the copy number of maltose metabolism and transport genes *(IMA1, IMA5, MAL31,* and *YPR196W-* 2 copies of each gene), flocculation related gene *(FLO1-* 2 copies) and hexose transporter *(HXT8-* 3 copies). Increased dosage of these genes in *S. jurei* could have conferred selective advantage towards better sugar utilization (Adamczyk et al. 2016; Lin and Li 2011; Ozcan and Johnston 1999; Soares 2011). Genes encoding for PAU proteins (a member of the seripauperin multigene family), copper resistance and salt tolerance related genes were found to be present in fewer copies in *S. jurei* genome compared to *S. cerevisiae.* This variation in copy number of genes in a genome can have phenotypic and physiological effects on the species (Adamo et al. 2012; Gorter de Vries et al. 2017; Landry et al. 2006).

We also searched for the presence of repetitive elements in *S. jurei* NCYC 3947^T^ and NCYC 3962 using BLAST and compared them to the Ty elements in *S. cerevisiae.* We detected Ty1-LTR, Ty2-LTR, Ty2-I-int, Ty3-LTR, Ty3-I and Ty4 sequences in both strains of *S. jurei.* Interestingly, we found an increased number of TY1-LTR, TY2-LTR, TY3-LTR and TY4 elements in *S. jurei* genome compared to *S. cerevisiae* (Table 7). Repetitive sequences are found in genomes of all eukaryotes and can be a potential source of genomic instability since they can recombine and cause chromosomal rearrangements, such as translocations, inversions and deletions (Chan and Kolodner 2011; Naseeb et al. 2016; Shibata et al. 2009).

**Table 7:** Counts of Ty elements in *S. cerevisiae, S. jurei* NCYC 3947^T^ and NCYC 3962.

| **Ty elements** | **Ty elements annotation** | **Counts in *S. cerevisiae*** | **Counts in *S. jurei* NCYC 3947<sup>T</sup>** | **Counts in *S. jurei* NCYC 3962** |
| --- | --- | --- | --- | --- |
| Ty | Yeast Ty transposable element Ty-pY109 near tRNA-Lys1 gene | 164 | 71 | 74 |
| Ty1-LTR | Ty1 LTR-retrotransposon from yeast (LTR) | 124 | 276 | 272 |
| Ty2-LTR | Ty2 LTR-retrotransposon from yeast (LTR) | 108 | 118 | 117 |
| Ty2-I-int | Ty2 LTR-retrotransposon from yeast (internal portion). | 15 | 2 | 2 |
| Ty3-LTR | *S. paradoxus* Ty3-like retrotransposon, Long terminal repeat | 61 | 70 | 71 |
| Ty3-I | *S. paradoxus* Ty3-like retrotransposon, Internal region. | 2 | 1 | 1 |
| Ty4 | Gag homolog, Ty4B=protease, integrase, reverse transcriptase, and RNase H domain containing protein {retrotransposon Ty4} | 51 | 164 | 162 |

### *Saccharomyces jurei* share a chromosomal translocation with *Saccharomyces mikatae* IFO 1815

To check the presence or absence of genomic rearrangements in *S. jurei,* we compared the chromosome structures between *S. jurei* NCYC 3947^T^ and *S. jurei* NCYC 3962 (Figure 1A), between *S. cerevisiae* S288C and *S. mikatae* IFO1815 (Figure 1B), between *S. jurei* NCYC 3947^T^ and *S. cerevisiae* S288C (Figure 2A) and between *S. jurei* NCYC 3947^T^ and *S. mikatae* IFO1815 (Figure 2B). The two *S. jurei* strains had a syntenic genome (Figure 1A), while we identified two chromosomal translocations with *S. cerevisiae* S288C (Figure 2A). One translocation is unique to *S. jurei* and is located between chromosomes I and XIII (Figure 2, red ovals), while the second translocation is located between chromosomes VI and VII in the same position of the previously identified translocation in *S. mikatae* IFO1815 (Figure 2, black ovals).

**Figure 1:**
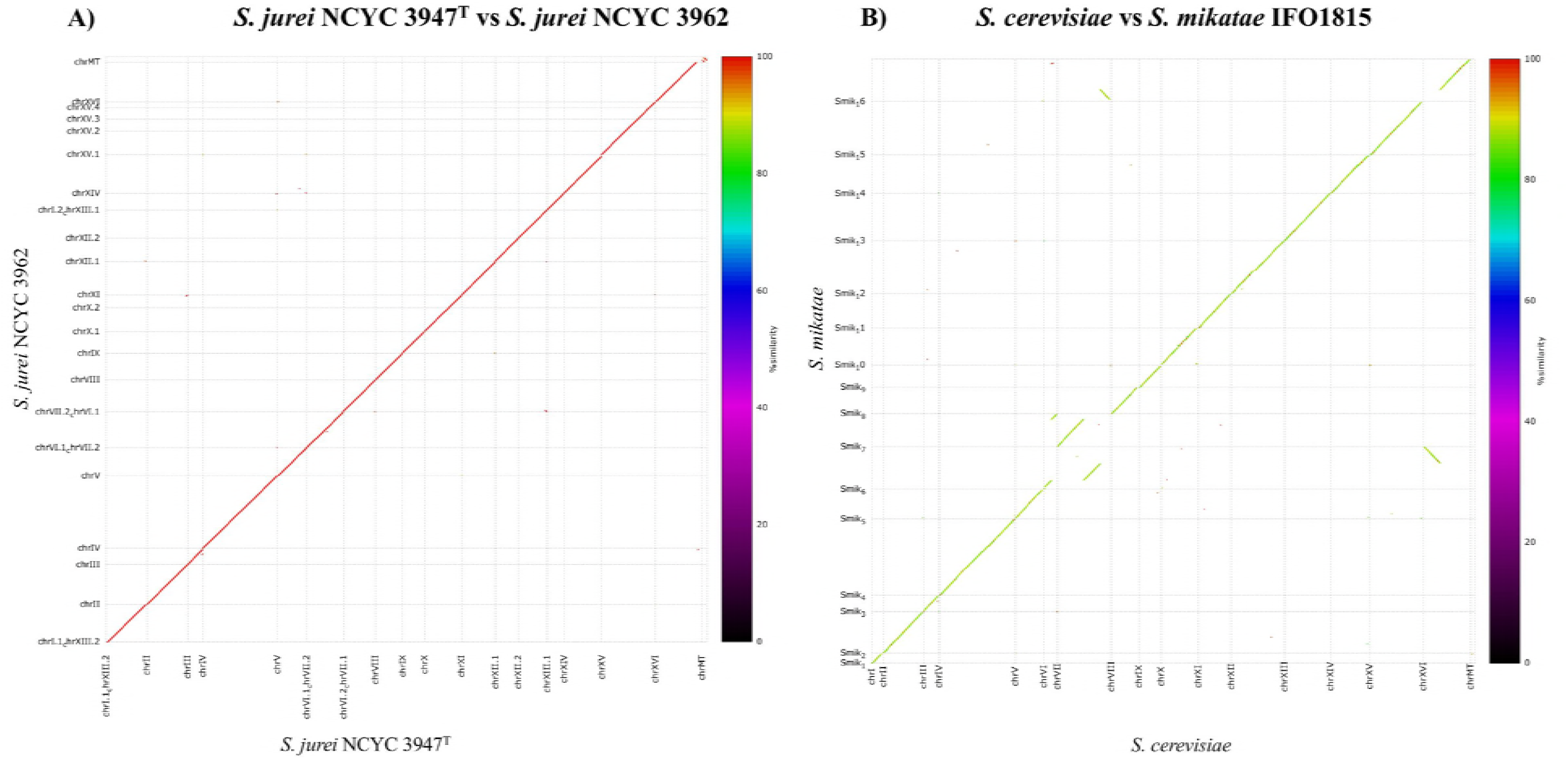
Dot plot alignments comparing the chromosome sequence identity of *S. jurei* NCYC 3947^T^ versus *S. jurei* NCYC 3962 (**A**) and *S. cerevisiae* S288C versus *S. mikatae* IFO1815 (**B**). The broken lines represent chromosomal translocations between chromosomes VI / VII and XVI / VII.

**Figure 2:**
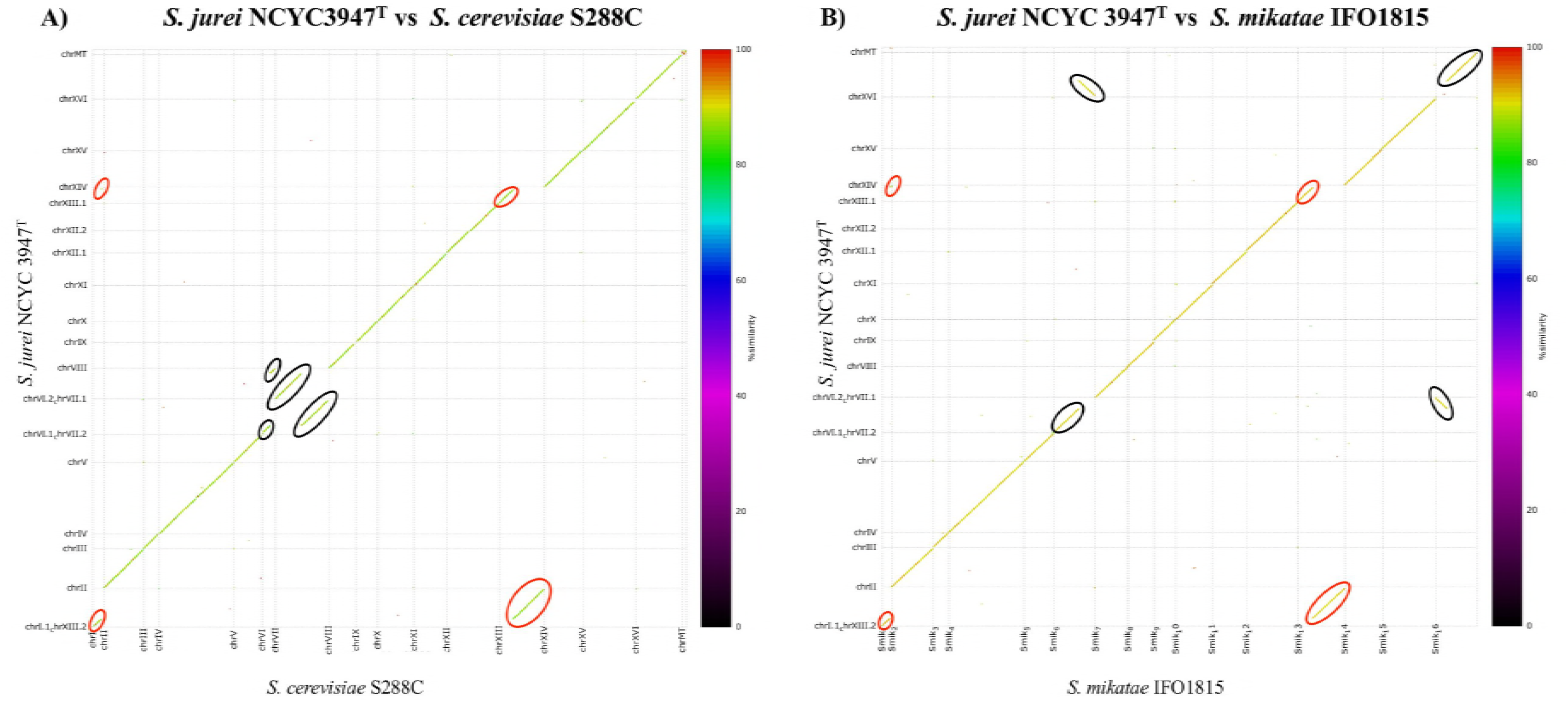
Dot plot alignments comparing the chromosome sequence identity of *S. jurei* NCYC 3947^T^ versus *S. cerevisiae* S288C (**A**) and *S. jurei* NCYC 3947^T^ versus *S. mikatae* IFO1815 (**B**). Black ovals represent the translocation between chromosomes VI and VII, which is common in *S. mikatae* and *S. jurei* whereas red ovals represent the translocation between chromosomes I and XIII, which is unique to *S. jurei.*

The breakpoints of the translocation I/XIII and are in the intergenic regions between uncharacterized genes. The breakpoints neighborhood is surrounded by several Ty elements (Ty1-LTR, Ty4, and Ty2-LTR) and one tRNA, which may have caused the rearrangement (Bridier-Nahmias et al. 2015; Fischer et al. 2000; Liti et al. 2013; Mieczkowski et al. 2006). The translocation in common with *S. mikatae* shares the same breakpoints between open reading frames (ORFs) YFR006w and YFR009w on chromosome VI, and between ORFs YGR021w and YGR026w on chromosome VII. This translocation is also shared by both strains of *S. mikatae*, but not with other *Saccharomyces sensu stricto* species. Overall this suggests a common evolutionary history between these stains and species, however an adaptive value of this rearrangement or a case of breakpoint re-usage cannot be ruled out since rearrangements can be adaptive with evidence both from nature and lab setting. (Adams et al. 1992; Avelar et al. 2013; Chang et al. 2013; Colson et al. 2004; Dunham et al. 2002; Fraser et al. 2005; Hewitt et al. 2014). Several natural isolates of *S. cerevisiae* present karyotypic changes (Hou et al. 2014) and the reciprocal translocation present between chromosomes VIII and XVI is able to confer sulphite resistance to the yeasts strains in vineyards (Perez-Ortin et al. 2002). Furthermore, lab experimental evolution studies in different strains of *S. cerevisiae* when evolved under similar condition end up sharing the same breakpoints (Dunham et al. 2002). Previous studies on mammalian systems have shown that breakpoints maybe reused throughout evolution at variable rates (Larkin et al. 2009; Murphy et al. 2005), and breakpoint re-usage has also been found between different strains of *S. pastorianus* (Hewitt et al. 2014).

### Novel genes present in *S. jurei*

The comparison between *S. jurei* and *S. cerevisiae* genome showed 622 differentially present genes. 179 open reading frames (ORFs) were predicted to be novel in *S. jurei* when compared to *S. cerevisiae* reference S288C strain (Table S5). To further confirm if these ORFs were truly novel, we analyzed the sequences in NCBI nucleotide database and in Saccharomyces Genome Database (SGD) against all the fungal species. We found 4 novel ORFs that have no significant match to any of the available genomes (Table S5-shown in red). 5 ORFs gave partial similarity to different fungal species such as *Vanderwaltozyma polyspora*, *Kluyveromyces marxianus*, *Torulaspora delbrueckii*, *Zygosaccharomyces rouxii*, *Hyphopichia burtonii, Kazachstania africana, Trichocera brevicornis, Lachancea walti,* and *Naumovozyma castellii* (Table S5-yellow highlighted). Majority of the remaining sequences gave full or partial matches to *S. cerevisiae* natural isolates (Strope et al. 2015), *S. paradoxus, S. mikatae, S. kudriavzevii, S. bayanus, S. uvarum,* and *S. eubayanus.*

Moreover, we also found 462 ORFs, which are present in *S. cerevisiae* genome but were lost in *S. jurei* (Table S6). The majority of genes which were novel or lost in *S. jurei* were found to be subtelomeric or telomeric, in regions known to show higher genetic variations (Bergstrom et al. 2014).

The genes lost in *S. jurei* encompass functionally verified ORFs, putative genes and uncharacterized genes. Some of the verified ORFs included ribosomal subunits genes, asparagine catabolism genes, alcohol dehydrogenase genes, hexose transporters, genes involved in providing resistance to arsenic compounds, phosphopyruvate hydratase genes, iron transport facilitators, ferric reductase genes and flocculation related genes.

We found that *S. jurei* genome lacks four out of seven alcohol dehydrogenase (AAD) genes including the functional *AAD4* gene, which is involved in oxidative stress response (Delneri et al. 1999a; Delneri et al. 1999b). Although *S. jurei* has lost *AAD4* gene, however, it was able to tolerate oxidative stress caused by 4mM H_2_O_2_ (Figure 3A).

**Figure 3:**
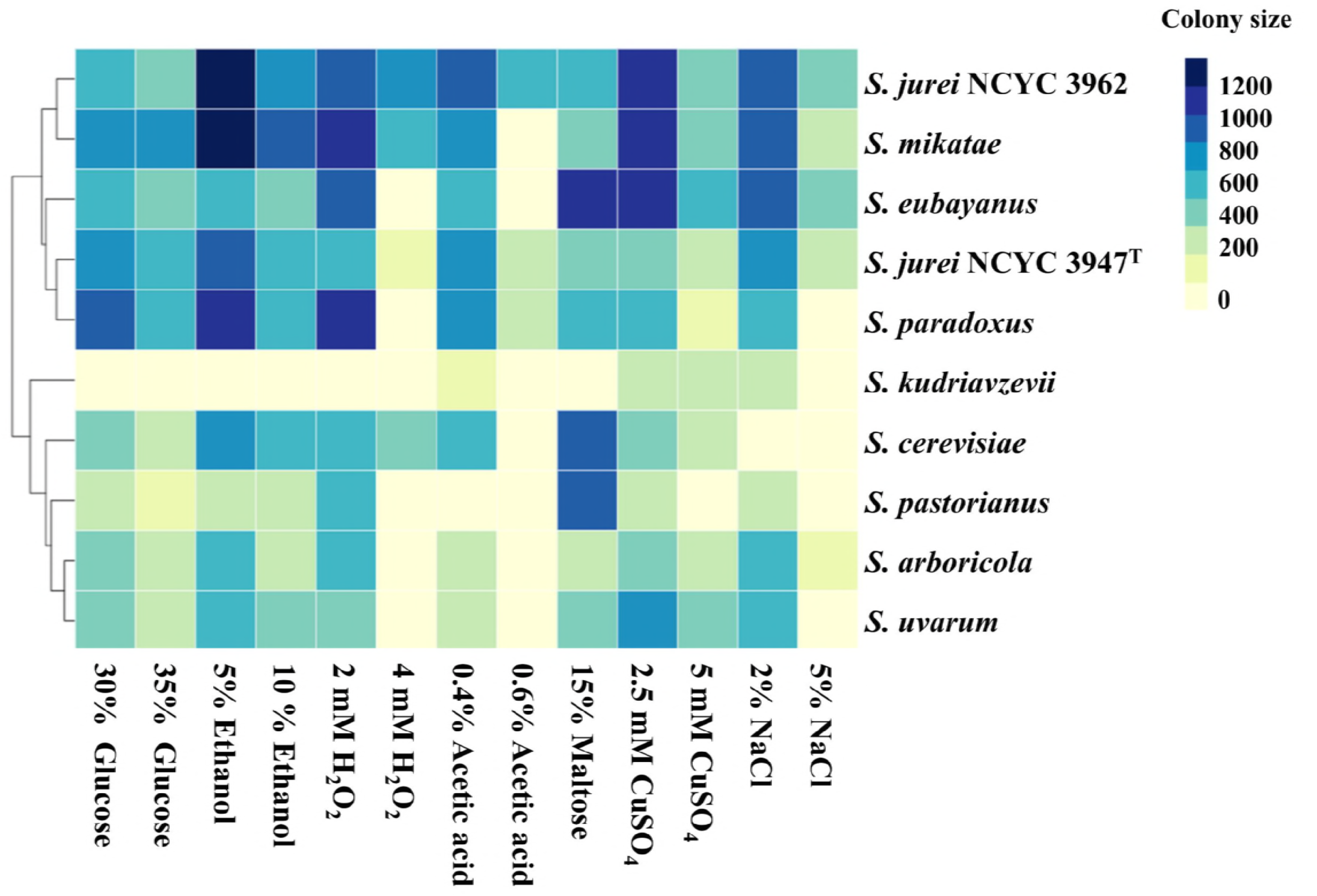
Heat map representing phenotypic fitness of *S. jurei* NCYC 3947^T^ and NCYC 3962 compared to *sensu stricto* species type strains in response to different environmental stressors at 30°C. Phenotypes are represented with colony sizes calculated as pixels and coloured according to the scale, with light yellow and dark blue colours representing the lowest and highest growth respectively. Hierarchical clustering of the strains is based on the overall growth profile under different media conditions tested.

All four genes of the *ASP3* gene cluster located on chromosome XII are absent in *S. jurei*. It was not surprising since this gene cluster is only known to be present in *S. cerevisiae* strains isolated from industrial and laboratory environments and lost from 128 diverse fungal species (Gordon et al. 2009; League et al. 2012). These genes are up-regulated during nitrogen starvation allowing the cells to grow by utilizing extracellular asparagine as a nitrogen source.

The hexose transporter family consists of 20 putative HXT genes *(HXT1-HXT17, GAL2, SNF3,* and *RGT2)* located on different chromosomes (Boles and Hollenberg 1997; Kruckeberg 1996) of which *HXT15, HXT16* and *HXT2* are absent from *S. jurei.* Under normal conditions, only 6 HXT genes *(HXT1* and *HXT3-HXT7)* are known to play role in glucose uptake suggesting that loss of 3 HXT genes from *S. jurei* is unlikely to affect glucose transport (Lin and Li 2011).

### Heterozygosis and strain divergence in the *S. jurei*

To detect genetic divergence between the two strains we mapped SNPs between the strains (NCYC 3947^T^ vs NCYC 3962), while to detect heterozygosis, we mapped the Single Nucleotide Polymorphisms (SNPs) in the two sets of alleles within the novel strains (NCYC 3947^T^ vs NCYC 3947^T^, and NCYC 3962 vs NCYC 3962). We found 6227 SNPs between the two strains, showing a genetic divergence between them. Moreover, 278 and 245 SNPs were found within NCYC 3947^T^ and NCYC 3962 strains respectively, indicating a low level of heterozygosity within each strain (Table 8). 139 SNPs were found be to common to both strains. Previous studies on *S. cerevisiae* and *S. paradoxus* strains from different lineages have shown that the level of heterozygosity is variable, with a large number of strains showing high level of heterozygosity isolated from human associated environments (Magwene et al. 2011; Tsai et al. 2008). A more recent study on 1011 *S. cerevisiae* natural strains showed that 63% of the sequenced isolates were heterozygous (Peter et al. 2018a).

**Table 8:** SNPs count in *S. jurei* NCYC 3947^T^ and NCYC 3962 genome.

| **Reference genome** | **Genome mapped** | **SNPs** |
| --- | --- | --- |
| NCYC 3947<sup>T</sup> | NCYC 3947<sup>T</sup> | 278 |
| NCYC 3962 | NCYC 3962 | 245 |
| NCYC 3947<sup>T</sup> | NCYC 3962 | 5702 |
| NCYC 3962 | NCYC 3947<sup>T</sup> | 6227 |

### Phylogenetic analysis

A first phylogeny construction using ITS/D1+D2 sequence analysis showed that *S. jurei* is placed in the tree close to *S. cerevisiae, S. mikatae* and *S. paradoxus* (Naseeb et al. 2017b). Here, we reconstructed the phylogeny using a multigene concatenation approach, which combines many genes together giving a large alignment (Baldauf et al. 2000; Brown et al. 2001; Fitzpatrick et al. 2006). Combination of concatenated genes improves the phylogenetic accuracy and helps to resolve the nodes and basal branching (Rokas et al. 2003). To reconstruct the evolutionary events, we concatenated 101 universally distributed orthologs obtained from complete genome sequencing data (Table S7). Both novel strains were located in one single monophyletic group, with the *S. mikatae* (Figure 4). Since *S. jurei* also have a chromosomal translocation in common with *S. mikatae*, it further shows that the two species share similar evolutionary history and hence present in the same group on the phylogenetic tree.

**Figure 4:**
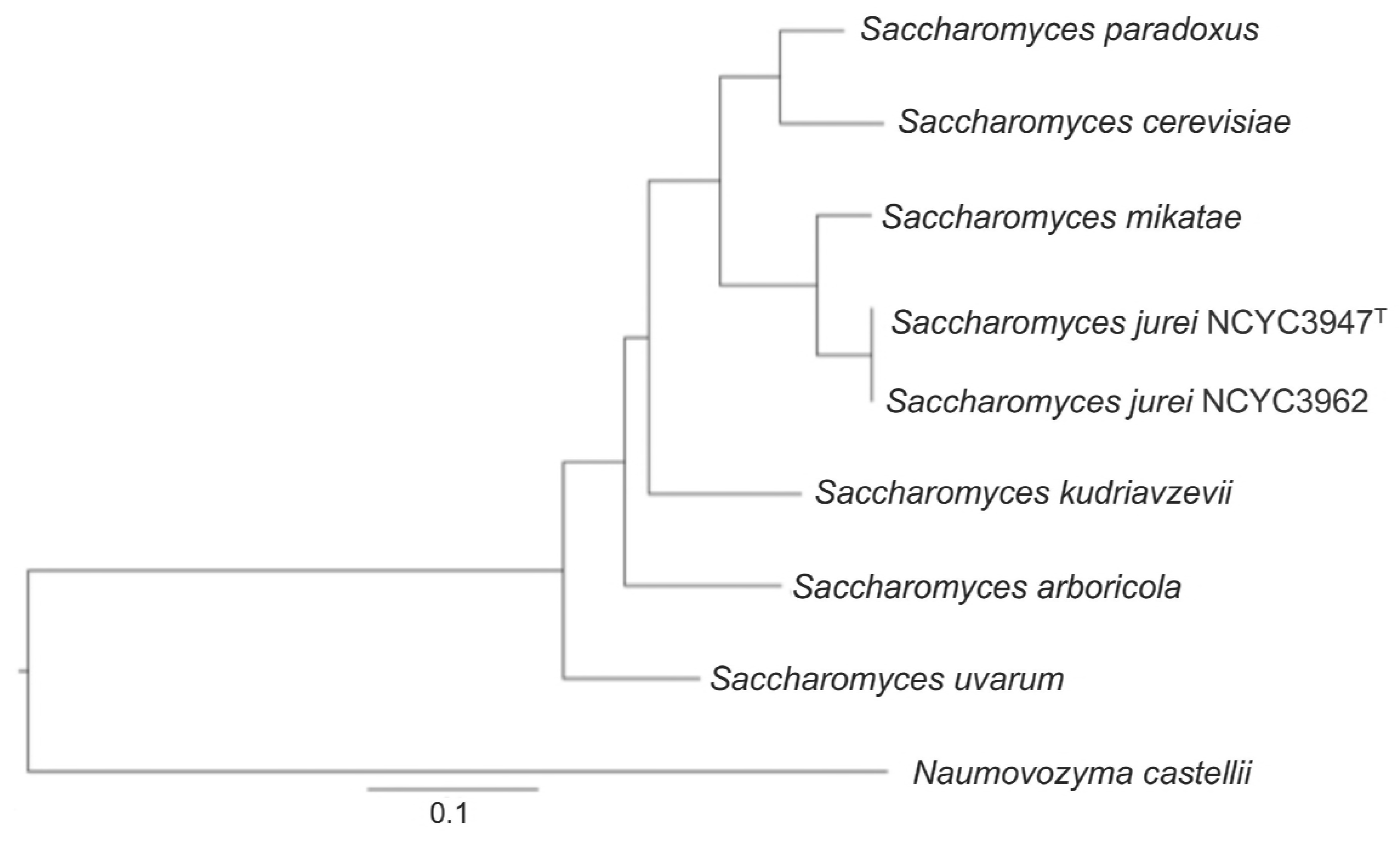
Phylogenetic tree showing both novel strains located in one single monophyletic group, with the *S. mikatae.* Maximum likelihood phylogeny was constructed using a concatenated alignment of 101 universally distributed genes. Sequences from all *Saccharomyces sensu stricto* species were aligned using StatAlign v3.1 and phylogenetic tree was built using RaxML 8.1.3 with *N. castellii* kept as out-group.

### Introgression analysis

To determine whether the two *S. jurei* strains possessed any introgressed region from other yeast species, we compared *S. jurei* genome with those of *S. cerevisiae*, *S. mikatae*, *S. paradoxus* and *S. kudriavzevii*. We did not observe introgression of any full-length genes or large segments of the genome (>1000 bp) in *S. jurei.* However, in both novel strains, we identified seven small DNA fragments (300 bp-700 bp) belonging to five different genes, which may have derived from *S. paradoxus* or *S. mikatae* (Table S8). DNA fragments from all the genes *(CSS3, IMA5, MAL33, YAL003W)* with the exception of *YDR541C,* showed a high sequence similarity to *S. paradoxus* genome, indicating putative introgression from *S. paradoxus* to *S. jurei* (Table S8).

Introgression of genetic material can easily occur in *Saccharomyces* species by crossing the isolates to make intraspecific or interspecific hybrids (Fischer et al. 2000; Naumov et al. 2000). Among the *Saccharomyces sensu stricto* group, introgressions have been demonstrated in natural and clinical yeast isolates (Liti et al. 2006; Muller and McCusker 2009; Wei et al. 2007; Zhang et al. 2010) and in wine, beer and other fermentation environments (de Barros Lopes et al. 2002; Dunn et al. 2012; Usher and Bond 2009). It is generally believed that introgressed regions are retained, as they may be evolutionarily advantageous (Strope et al. 2015; Novo et al. 2009). Previous studies have demonstrated that introgression in *S. cerevisiae* is relatively common and a majority of the genes are derived from introgression with *S. paradoxus* (Liti et al. 2006; Novo et al. 2009; Peter et al. 2018a; Strope et al. 2015; Warringer et al. 2011).

### Phenotypic profiling of *S. jurei*

We performed large-scale phenotypic profiling under various stress conditions and at different temperatures to capture the fitness landscape of *S. jurei* (strains NCYC 3947^T^ and NCYC 3962) relative to other *Saccharomyces sensu stricto* species. Colony size was taken as a proxy for fitness score (see methods). Generally the fitness of *S. jurei* NCYC 3962 in different environmental stressor conditions was higher compared to *S. jurei* NCYC 3947^T^ (Figure 3). Remarkably, only *S. jurei* NCYC 3962 was able to grow well on higher concentrations of acetic acid (Figure 3). Like most of the other *Saccharomyces* yeast species, both strains of *S. jurei* can also grow in media containing 10% ethanol. Although *S. eubayanus* showed the highest fitness in media containing 15% maltose, both strains of *S. jurei* were also able to tolerate high concentrations of maltose. Moreover, *S. jurei* NCYC 3962 was able to better tolerate higher concentrations of H_2_O_2_, CuSO_4_ and NaCl compared to most of the other *sensu stricto* species (Figure 3). *Saccharomyces* yeast species can acquire copper tolerance either due to an increase in *CUP1* copy number (Warringer et al. 2011) or due to the use of copper sulfate as a fungicide in vineyards (Fay et al. 2004; Perez-Ortin et al. 2002). The genomic analysis shows that both strains of *S. jurei* possess one copy of *CUP1*, indicating other factors maybe associated with copper tolerance.

Phenotypically, both strains of *S. jurei* clustered with *S. mikatae* and *S. paradoxus,* which is in accordance with our phylogenetic results, and, interestingly, the brewing yeast *S. eubayanus* was also present in the same cluster (Figure 3). This may indicate that in spite of the phylogenetic distance, *S. eubayanus* may have shared similar ecological conditions with the other above mentioned species.

We also evaluated the fitness of *S. jurei* strains in comparison to *Saccharomyces sensu stricto* species at different temperatures, taking into account growth parameters such as lag phase (λ), maximum growth rate (μ_max_), and maximum biomass (*A*_max_) (Tables S9-S11). The optimum growth of NCYC 3947^T^ and NCYC 3962 was at 25°C and 30°C respectively (Table S10). Both strains of *S. jurei* are able to grow at a high temperatures (i.e. 37°C) compared to *S. kudriavzevii, S. pastorianus, S. arboricola, S. uvarum,* and *S. eubayanus,* which are unable to grow at 37° C (Table S10). The ability of *S. jurei* strains to grow well both at cold and warm suggest that this species evolved to be a generalist rather than a specialist in terms of thermoprofiles. The growth profiles captured at different temperatures for the other *Saccharomyces* species was in accordance to the previously published study (Salvado et al. 2011).

## Conclusions

High quality *de novo* assembly of two novel strains of *S. jurei* (NCYC 3947^T^ and NCYC 3962) has been carried out using short and long reads sequencing strategies. We obtained a 12Mb genome and were able to assemble full chromosomes of both species. We found two reciprocal chromosomal translocations in *S. jurei* genome, between chromosomes I/XIII and VI/VII. The translocation between chromosomes I/XIII is unique to *S. jurei* genome, whereas the translocation between VI/VII is shared with *S. mikatae* IFO1815 and IFO1816. This suggests a common origin between *S. jurei* and *S. mikatae* and *S. jurei* evolved after acquiring the translocation between chromosomes I/XIII, while *S. mikatae* 1815 acquired a second translocation between chromosomes XVI/VII. Moreover, both strains of *S. jurei* showed low heterozygosis within themselves and were genetically diverged possessing 6227 SNPs between them. We found 4 novel ORFs that had no significant match to any of the available genomes. *S. jurei* genome had an increased number of TY elements compared to *S. cerevisiae* and showed no signatures of introgression. The phylogenetic analysis showed that the novel species is closely related to *S. mikatae,* forming a single monophyletic group.

Phenotypically, the environmental stressor profiles of *S. jurei* are similar to those of with *S. mikatae*, *S. paradoxus*, *S. cerevisiae* (which further reiterate that *S. jurei* is closely related to these species) and *S. eubayanus.* We found that *S. jurei* NCYC 3962 compared to other *sensu stricto* species was able to grow well at high concentrations of acetic acid. In general, *S. jurei* NCYC 3962 showed relatively higher fitness compared to *S. jurei* NCYC 3947^T^ under most of the environmental stress conditions tested. Both strains of *S. jurei* showed similar growth rate at relatively low temperature, however, NCYC 3962 showed increased fitness compared to NCYC 3947^T^ at higher temperatures. The sequencing data and the large-scale phenotypic screening of this new species provide the basis for future investigations of biotechnological and industrial importance.

## Data accessibility

The sequences and annotations reported in this paper are available in the European Nucleotide Archive under project ID PRJEB24816, assembly ID GCA_900290405 and accession number ERZ491603.

## Competing interests

The authors declare no competing interests.

## Acknowledgements

The authors would like to thank Genomic Technologies Core Facility at the University of Manchester for Illumina Hi-seq and Dr. Haiping Hao at Deep Sequencing and Microarray Core Facility of Johns Hopkins University for PacBio sequencing. SN is supported through BBSRC funding (BB/L021471/1). HA is supported by a scholarship funded by the Kuwait government through Kuwait University.

## Supplementary Figure Legend

**Figure S1**: Alignment of the amino acid sequences of *MEL1* belonging to *S. jurei* NCYC 3947^T^ (Sj) and *S. mikatae* IFO 1816 (Sm) (NCBI Accession number: Q11129.1) *MEL1* genes. Sequences were aligned using Clustal Omega. Asterisk (*) indicate a position of conserved residue, a colon (:) indicate a position of strong conservation between the alignment, and a period (.) indicate a position of weak conservation between the alignment.

**Supplementary Table Legends**

**Table S1**: List of genes which are present in simple one to one orthologous relationship.

**Table S2**: List of genes which are present in many to many relationship.

**Table S3**: List of genes which are present in many to one relationship.

**Table S4**: List of genes which are present in one to many relationship.

**Table S5**: List of genes which are present in *S. cerevisiae* but absent in *S. jurei.*

**Table S6**: List of genes which are present in *S. jurei* but absent in *S. cerevisiae.*

**Table S7**: List of genes which are used to construct the phylogenetic tree.

**Table S8**: List of genes which are potentially introgressed in *S. jurei* genome from *S. paradoxus*.

**Table S9**: Lag phase time (A) of *Saccharomyces species* used in this study at different temperatures.

**Table S10**: Maximum growth rate (μmax) of *Saccharomyces* species used in this study at different temperatures.

**Table S11**: Maximum biomass (A_max_) of *Saccharomyces species* used in this study at different temperatures.

## References

Adamczyk, J., et al. (2016), ‘Copy number variations of genes involved in stress responses reflect the redox state and DNA damage in brewing yeasts’, Cell Stress Chaperones, 21 (5), 849–64.

Adamo, G. M., et al. (2012), ‘Amplification of the CUP1 gene is associated with evolution of copper tolerance in Saccharomyces cerevisiae’, Microbiology, 158 (Pt 9), 2325–35.

Adams, J., et al. (1992), ‘Adaptation and major chromosomal changes in populations of Saccharomyces cerevisiae’, Curr Genet, 22 (1), 13–9.

Avelar, A. T., et al. (2013), ‘Genome architecture is a selectable trait that can be maintained by antagonistic pleiotropy’, Nat Commun, 4, 2235.

Baker, E., et al. (2015), ‘The Genome Sequence of Saccharomyces eubayanus and the Domestication of Lager-Brewing Yeasts’, Mol BiolEvol, 32 (11), 2818–31.

Baldauf, S. L., et al. (2000), ‘A kingdom-level phylogeny of eukaryotes based on combined protein data’, Science, 290 (5493), 972–7.

Bao, W., Kojima, K. K., and Kohany, O. (2015), ‘Repbase Update, a database of repetitive elements in eukaryotic genomes’, Mob DNA, 6, 11.

Berglund, A. C., et al. (2008), ‘InParanoid 6: eukaryotic ortholog clusters with inparalogs’, Nucleic Acids Res, 36 (Database issue), D263–6.

Bergstrom, A., et al. (2014), ‘A high-definition view of functional genetic variation from natural yeast genomes’, Mol Biol Evol, 31 (4), 872–88.

Boles, E. and Hollenberg, C. P. (1997), ‘The molecular genetics of hexose transport in yeasts’, FEMS Microbiol Rev, 21 (1), 85–111.

Bourque, G., et al. (2005), ‘Comparative architectures of mammalian and chicken genomes reveal highly variable rates of genomic rearrangements across different lineages’, Genome Res, 15 (1), 98–110.

Brice, C., et al. (2018), ‘Adaptability of the Saccharomyces cerevisiae yeasts to wine fermentation conditions relies on their strong ability to consume nitrogen’, PLoS One, 13 (2), e0192383.

Bridier-Nahmias, A., et al. (2015), ‘Retrotransposons. An RNA polymerase III subunit determines sites of retrotransposon integration’, Science, 348 (6234), 585–8.

Brown, J. R., et al. (2001), ‘Universal trees based on large combined protein sequence data sets’, Nat Genet, 28 (3), 281–5.

Byrne, K. P. and Wolfe, K. H. (2005), ‘The Yeast Gene Order Browser: combining curated homology and syntenic context reveals gene fate in polyploid species’, Genome Res, 15 (10), 1456–61.

Cardinali, G. and Martini, A. (1994), ‘Electrophoretic karyotypes of authentic strains of the sensu stricto group of the genus Saccharomyces’, Int JSyst Bacteriol, 44 (4), 791–7.

Carle, G. F. and Olson, M. V. (1985), ‘An electrophoretic karyotype for yeast’, Proc Natl AcadSci USA, 82 (11), 3756–60.

Casaregola, S., et al. (2000), ‘Genomic exploration of the hemiascomycetous yeasts: 17. Yarrowia lipolytica’, FEBS Lett, 487 (1), 95–100.

Challis, D., et al. (2012), ‘An integrative variant analysis suite for whole exome next-generation sequencing data’, BMC Bioinformatics, 13, 8.

Chan, J. E. and Kolodner, R. D. (2011), ‘A genetic and structural study of genome rearrangements mediated by high copy repeat Ty1 elements’, PLoS Genet, 7 (5), e1002089.

Chang, S. L., et al. (2013), ‘Dynamic large-scale chromosomal rearrangements fuel rapid adaptation in yeast populations’, PLoS Genet, 9 (1), e1003232.

Chin, C. S., et al. (2013), ‘Nonhybrid, finished microbial genome assemblies from long-read SMRT sequencing data’, Nat Methods, 10 (6), 563–9.

Cliften, P., et al. (2003), ‘Finding functional features in Saccharomyces genomes by phylogenetic footprinting’, Science, 301 (5629), 71–6.

Colson, I., Delneri, D., and Oliver, S. G. (2004), ‘Effects of reciprocal chromosomal translocations on the fitness of Saccharomyces cerevisiae’, EMBO Rep, 5 (4), 392–8.

de Barros Lopes, M., et al. (2002), ‘Evidence for multiple interspecific hybridization in Saccharomyces sensu stricto species’, FEMS Yeast Res, 1 (4), 323–31.

Delneri, D., Gardner, D. C., and Oliver, S. G. (1999a), ‘Analysis of the seven-member AAD gene set demonstrates that genetic redundancy in yeast may be more apparent than real’, Genetics, 153 (4), 1591–600.

Delneri, D., et al. (1999b), ‘Disruption of seven hypothetical aryl alcohol dehydrogenase genes from Saccharomyces cerevisiae and construction of a multiple knock-out strain’, Yeast, 15 (15), 1681–9.

Dunham, M. J., et al. (2002), ‘Characteristic genome rearrangements in experimental evolution of Saccharomyces cerevisiae’, Proc Natl Acad Sci U S A, 99 (25), 16144–9.

Dunn, B., et al. (2012), ‘Analysis of the Saccharomyces cerevisiae pan-genome reveals a pool of copy number variants distributed in diverse yeast strains from differing industrial environments’, Genome Res, 22 (5), 908–24.

Engel, S. R. and Cherry, J. M. (2013), ‘The new modern era of yeast genomics: community sequencing and the resulting annotation of multiple Saccharomyces cerevisiae strains at the Saccharomyces Genome Database’, Database-the Journal of Biological Databases and Curation.

Fay, J. C., et al. (2004), ‘Population genetic variation in gene expression is associated with phenotypic variation in Saccharomyces cerevisiae’, Genome Biol, 5 (4), R26.

Fischer, G., et al. (2000), ‘Chromosomal evolution in Saccharomyces’, Nature, 405 (6785), 451–4.

Fischer, G., et al. (2001), ‘Evolution of gene order in the genomes of two related yeast species’, Genome Res, 11 (12), 2009–19.

Fischer, G., et al. (2006), ‘Highly variable rates of genome rearrangements between hemiascomycetous yeast lineages’, PLoS Genet, 2 (3), e32.

Fitzpatrick, D. A., et al. (2006), ‘A fungal phylogeny based on 42 complete genomes derived from supertree and combined gene analysis’, BMC Evol Biol, 6, 99.

Fraser, J. A., et al. (2005), ‘Chromosomal translocation and segmental duplication in Cryptococcus neoformans’, Eukaryot Cell, 4 (2), 401–6.

Fujita, S. and Hashimoto, T. (2000), ‘DNA fingerprinting patterns of Candida species using HinfI endonuclease’, Int JSystEvolMicrobiol, 50 Pt 3, 1381–9.

Gertz, E. M., et al. (2006), ‘Composition-based statistics and translated nucleotide searches: improving the TBLASTN module of BLAST’, BMC Biol, 4, 41.

Goddard, M. R. and Greig, D. (2015), ‘Saccharomyces cerevisiae: a nomadic yeast with no niche?’, FEMS Yeast Res, 15 (3).

Goffeau, A., et al. (1996), ‘Life with 6000 genes’, Science, 274 (5287), 546, 63–7.

Gordon, J. L., Byrne, K. P., and Wolfe, K. H. (2009), ‘Additions, losses, and rearrangements on the evolutionary route from a reconstructed ancestor to the modern Saccharomyces cerevisiae genome’, PLoS Genet, 5 (5), e1000485.

Gorter de Vries, A. R., Pronk, J. T., and Daran, J. G. (2017), ‘Industrial Relevance of Chromosomal Copy Number Variation in Saccharomyces Yeasts’, Appl Environ Microbiol, 83 (11).

Hall, C., Brachat, S., and Dietrich, F. S. (2005), ‘Contribution of horizontal gene transfer to the evolution of Saccharomyces cerevisiae’, Eukaryot Cell, 4 (6), 1102–15.

Hewitt, S. K., et al. (2014), ‘Sequencing and characterisation of rearrangements in three S. pastorianus strains reveals the presence of chimeric genes and gives evidence of breakpoint reuse’, PLoS One, 9 (3), e92203.

Hou, J., et al. (2014), ‘Chromosomal rearrangements as a major mechanism in the onset of reproductive isolation in Saccharomyces cerevisiae’, Curr Biol, 24 (10), 1153–9.

Jones, P., et al. (2014), ‘InterProScan 5: genome-scale protein function classification’, Bioinformatics, 30 (9), 1236–40.

Jouhten, P., et al. (2016), ‘Saccharomyces cerevisiae metabolism in ecological context’, FEMS Yeast Res, 16 (7).

Kellis, M., et al. (2003), ‘Sequencing and comparison of yeast species to identify genes and regulatory elements’, Nature, 423 (6937), 241–54.

Kent, W. J., et al. (2002), ‘The human genome browser at UCSC’, Genome Res, 12 (6), 996–1006.

Kruckeberg, A. L. (1996), ‘The hexose transporter family of Saccharomyces cerevisiae’, Arch Microbiol, 166 (5), 283–92.

Kurtz, S., et al. (2004), ‘Versatile and open software for comparing large genomes’, Genome Biol, 5 (2), R12.

Landry, C. R., et al. (2006), ‘Genome-wide scan reveals that genetic variation for transcriptional plasticity in yeast is biased towards multi-copy and dispensable genes’, Gene, 366 (2), 343–51.

Langmead, B. and Salzberg, S. L. (2012), ‘Fast gapped-read alignment with Bowtie 2’, Nat Methods, 9 (4), 357–9.

Larkin, D. M., et al. (2009), ‘Breakpoint regions and homologous synteny blocks in chromosomes have different evolutionary histories’, Genome Res, 19 (5), 770–7.

League, G. P., Slot, J. C., and Rokas, A. (2012), ‘The ASP3 locus in Saccharomyces cerevisiae originated by horizontal gene transfer from Wickerhamomyces’, FEMS Yeast Res, 12 (7), 859–63.

Lewis, S. E., et al. (2002), ‘Apollo: a sequence annotation editor’, Genome Biol, 3 (12), RESEARCH0082.

Li, H., et al. (2009), ‘The Sequence Alignment/Map format and SAMtools’, Bioinformatics, 25 (16), 2078–9.

Libkind, D., et al. (2011), ‘Microbe domestication and the identification of the wild genetic stock of lager-brewing yeast’, Proc Natl Acad Sci U S A, 108 (35), 14539–44.

Lin, Z. and Li, W. H. (2011), ‘Expansion of hexose transporter genes was associated with the evolution of aerobic fermentation in yeasts’, Mol Biol Evol, 28 (1), 131–42.

Liti, G., Barton, D. B., and Louis, E. J. (2006), ‘Sequence diversity, reproductive isolation and species concepts in Saccharomyces’, Genetics, 174 (2), 839–50.

Liti, G., et al. (2013), ‘High quality de novo sequencing and assembly of the Saccharomyces arboricolus genome’, BMC Genomics, 14, 69.

Liti, G., et al. (2009), ‘Population genomics of domestic and wild yeasts’, Nature, 458 (7236), 337–41.

Lowe, T. M. and Eddy, S. R. (1997), ‘tRNAscan-SE: a program for improved detection of transfer RNA genes in genomic sequence’, Nucleic Acids Res, 25 (5), 955–64.

Lynch, M. (2002), ‘Genomics. Gene duplication and evolution’, Science, 297 (5583), 945–7.

Magwene, P. M., et al. (2011), ‘Outcrossing, mitotic recombination, and life-history tradeoffs shape genome evolution in Saccharomyces cerevisiae’, Proc Natl Acad Sci U S A, 108 (5), 1987–92.

Martini, A. V. and Martini, A. (1987), ‘Three newly delimited species of Saccharomyces sensu stricto’, Antonie Van Leeuwenhoek, 53 (2), 77–84.

Masneuf, I., et al. (1998), ‘New hybrids between Saccharomyces sensu stricto yeast species found among wine and cider production strains’, Appl Environ Microbiol, 64 (10), 3887–92.

Mieczkowski, P. A., Lemoine, F. J., and Petes, T. D. (2006), ‘Recombination between retrotransposons as a source of chromosome rearrangements in the yeast Saccharomyces cerevisiae’, DNA Repair (Amst), 5 (9–10), 1010–20.

Muller, L. A. and McCusker, J. H. (2009), ‘A multispecies-based taxonomic microarray reveals interspecies hybridization and introgression in Saccharomyces cerevisiae’, FEMS Yeast Res, 9 (1), 143–52.

Murphy, W. J., et al. (2005), ‘Dynamics of mammalian chromosome evolution inferred from multispecies comparative maps’, Science, 309 (5734), 613–7.

Naseeb, S. and Delneri, D. (2012), ‘Impact of chromosomal inversions on the yeast DAL cluster’, PLoS One, 7 (8), e42022.

Naseeb, S., et al. (2017a), ‘Rapid functional and evolutionary changes follow gene duplication in yeast’, Proc Biol Sci, 284 (1861).

Naseeb, S., et al. (2016), ‘Widespread Impact of Chromosomal Inversions on Gene Expression Uncovers Robustness via Phenotypic Buffering’, Mol Biol Evol, 33 (7), 1679–96.

Naseeb, S., et al. (2017b), ‘Saccharomyces jurei sp. nov., isolation and genetic identification of a novel yeast species from Quercus robur’, Int J Syst Evol Microbiol, 67 (6), 2046–52.

Naumov, G. I. (1987), ‘Genetic basis for classification and identification of the ascomycetous yeasts.’, Stud. Mycol, 30, 469–75.

Naumov, G. I., Naumova, E. S., and Louis, E. J. (1995. b), ‘Two new genetically isolated populations of the Saccharomyces sensu stricto complex from Japan.’, J. Gen. App. Microbiol, 41, 499–505.

Naumov, G. I., Naumova, E. S., and Sancho, E. D. (1996), ‘Genetic reidentification of Saccharomyces strains associated with black knot disease of trees in Ontario and Drosophila species in California’, Can J Microbiol, 42, 335–39.

Naumov, G. I., et al. (1995.a), ‘A new genetically isolated population of the Saccharomyces sensu stricto complex from Brazil’, Antonie Van Leeuwenhoek, 67 (4), 351–5.

Naumov, G. I., et al. (2000), ‘Three new species in the Saccharomyces sensu stricto complex: Saccharomyces cariocanus, Saccharomyces kudriavzevii and Saccharomyces mikatae’, Int J Syst Evol Microbiol, 50 Pt 5, 1931–42.

Nawrocki, E. P. and Eddy, S. R. (2013), ‘Infernal 1.1: 100-fold faster RNA homology searches’, Bioinformatics, 29 (22), 2933–5.

Nawrocki, E. P., et al. (2015), ‘Rfam 12.0: updates to the RNA families database’, Nucleic Acids Res, 43 (Database issue), D130–7.

Nguyen, H. V., et al. (2011), ‘Deciphering the hybridisation history leading to the Lager lineage based on the mosaic genomes of Saccharomyces bayanus strains NBRC1948 and CBS380’, PLoS One, 6 (10), e25821.

Novo, M., et al. (2009), ‘Eukaryote-to-eukaryote gene transfer events revealed by the genome sequence of the wine yeast Saccharomyces cerevisiae EC1118’, Proc Natl Acad Sci U S A, 106 (38), 16333–8.

Ozcan, S. and Johnston, M. (1999), ‘Function and regulation of yeast hexose transporters’, Microbiol Mol Biol Rev, 63 (3), 554–69.

Perez-Ortin, J. E., et al. (2002), ‘Molecular characterization of a chromosomal rearrangement involved in the adaptive evolution of yeast strains’, Genome Res, 12 (10), 1533–9.

Peter, J., et al. (2018a), ‘Genome evolution across 1,011 Saccharomyces cerevisiae isolates’, Nature.

Peter, J., et al. (2018b), ‘Genome evolution across 1,011 Saccharomyces cerevisiae isolates’, Nature, 556 (7701), 339–44.

Querol, A. and Bond, U. (2009), ‘The complex and dynamic genomes of industrial yeasts’, FEMS Microbiol Lett, 293 (1), 1–10.

Raney, B. J., et al. (2014), ‘Track data hubs enable visualization of user-defined genome-wide annotations on the UCSC Genome Browser’, Bioinformatics, 30 (7), 1003–5.

Rokas, A., et al. (2003), ‘Genome-scale approaches to resolving incongruence in molecular phylogenies’, Nature, 425 (6960), 798–804.

Salvado, Z., et al. (2011), ‘Temperature adaptation markedly determines evolution within the genus Saccharomyces’, Appl Environ Microbiol, 77 (7), 2292–302.

Scannell, D. R., et al. (2011), ‘The Awesome Power of Yeast Evolutionary Genetics: New Genome Sequences and Strain Resources for the Saccharomyces sensu stricto Genus’, G3 (Bethesda), 1 (1), 11–25.

Seoighe, C., et al. (2000), ‘Prevalence of small inversions in yeast gene order evolution’, Proc Natl Acad Sci US A, 97 (26), 14433–7.

Shibata, Y., et al. (2009), ‘Yeast genome analysis identifies chromosomal translocation, gene conversion events and several sites of Ty element insertion’, Nucleic Acids Res, 37 (19), 6454–65.

Smit, A. F. A., Hubley, R., and Green, P. (2013–2015), ‘<**Error! Hyperlink reference not valid.**, RepeatMasker Open-4.0.

Soares, E. V. (2011), ‘Flocculation in Saccharomyces cerevisiae: a review’, J Appl Microbiol, 110 (1), 1–18.

Stanke, M. and Morgenstern, B. (2005), ‘AUGUSTUS: a web server for gene prediction in eukaryotes that allows user-defined constraints’, Nucleic Acids Res, 33 (Web Server issue), W465–7.

Strope, P. K., et al. (2015), ‘The 100-genomes strains, an S. cerevisiae resource that illuminates its natural phenotypic and genotypic variation and emergence as an opportunistic pathogen’, Genome Res, 25 (5), 762–74.

Tsai, I. J., et al. (2008), ‘Population genomics of the wild yeast Saccharomyces paradoxus: Quantifying the life cycle’, Proc Natl Acad Sci U S A, 105 (12), 4957–62.

Usher, J. and Bond, U. (2009), ‘Recombination between homoeologous chromosomes of lager yeasts leads to loss of function of the hybrid GPH1 gene’, Appl Environ Microbiol, 75 (13), 4573–9.

Walther, A., Hesselbart, A., and Wendland, J. (2014), ‘Genome sequence of Saccharomyces carlsbergensis, the world’s first pure culture lager yeast’, G3 (Bethesda), 4 (5), 783–93.

Wang, S. A. and Bai, F. Y. (2008), ‘Saccharomyces arboricolus sp. nov., a yeast species from tree bark’, Int J Syst Evol Microbiol, 58 (Pt 2), 510–4.

Warringer, J., et al. (2011), ‘Trait variation in yeast is defined by population history’, PLoS Genet, 7 (6), e1002111.

Wei, W., et al. (2007), ‘Genome sequencing and comparative analysis of Saccharomyces cerevisiae strain YJM789’, Proc Natl Acad Sci U S A, 104 (31), 12825–30.

Zhang, H., et al. (2010), ‘Saccharomyces paradoxus and Saccharomyces cerevisiae reside on oak trees in New Zealand: evidence for migration from Europe and interspecies hybrids’, FEMS Yeast Res, 10 (7), 941–7.

